# Sex-specific effects of intensity and dose of physical activity on BOLD-fMRI cerebrovascular reactivity and cerebral pulsatility

**DOI:** 10.1101/2024.10.10.617666

**Authors:** Zacharie Potvin-Jutras, Brittany Intzandt, Hanieh Mohammadi, Peiying Liu, Jean J. Chen, Claudine J. Gauthier

**Affiliations:** Department of Physics, Concordia University, Canada; School of Health, Concordia University, Canada; Centre ÉPIC, Montreal Heart Institute, Montréal, Québec, Canada; BrainLab, Hurvitz Brain Sciences Research Program, Sunnybrook Research Institute, Toronto, Ontario, Canada; Sandra Black Centre for Brain Resilience and Recovery, Sunnybrook Research Institute, Toronto, Ontario, Canada; Department of Medicine, Université de Montréal, Montréal, Québec, Canada; Department of Radiology, Johns Hopkins University School of Medicine, Baltimore, Maryland, USA; Department of Diagnostic Radiology & Nuclear Medicine, University of Maryland School of Medicine, Baltimore, Maryland, USA; Rotman Research Institute, Baycrest Academy for Research and Education, Toronto, Canada; Department of Medical Biophysics, University of Toronto, Toronto, Canada; Department of Biomedical Engineering, University of Toronto, Toronto, Canada

## Abstract

Cerebrovascular reactivity (CVR) and cerebral pulsatility (CP) are important indicators of cerebrovascular health and have been shown to be associated with physical activity (PA). Sex differences have been shown to influence the impact of PA on cerebrovascular health. However, the sex-specific effects of PA on CP and CVR, particularly in relation to intensity and dosage of PA, remains unknown. Thus, this cross-sectional study aimed to evaluate the sex-specific effects of different intensities and doses of PA on CVR and CP. The Human Connectome - Aging dataset was used, including 626 participants (350 females, 276 males) aged 36-85 (mean age: 58.8 ± 14.1 years). Females were stratified into premenopausal and postmenopausal groups to assess the potential influence of menopausal status. Novel tools based solely on resting state fMRI data were used to estimate both CVR and CP. The International Physical Activity Questionnaire was used to quantify weekly self-reported PA as metabolic equivalent of task. Results indicated that both sexes and menopausal subgroups revealed negative linear relationships between relative CVR and PA. Furthermore, females presented a unique non-linear relationship between relative CVR and total PA in the cerebral cortex. In females, there were also relationships with total and walking PA in occipital and cingulate regions. In males, we observed relationships between total or vigorous PA and CVR in parietal and cingulate regions. Sex-specific effects were also observed with CP, whereby females benefited across a greater number of regions and intensities than males, especially in the postmenopause group. Overall, males and females appear to benefit from different amounts and intensities of PA, with menopause status significantly influencing the effect of PA on cerebrovascular outcomes, underscoring the need for sex-specific recommendations in promoting cerebrovascular health.

## Introduction

Physical inactivity is associated with an increased risk of developing cardiovascular (Lippi et al., 2020) and neurological diseases (Kantawala et al., 2023). More specifically, physical inactivity is associated with greater arterial stiffness (Park et al., 2017), reduced endothelial function (Bowden Davies et al., 2021) and decreased vasodilatory function (Thijssen et al., 2010). Conversely, participating in physical activity (PA) has been shown to enhance vascular health, thereby delaying or preventing disease onset and progression (Green & Smith, 2018). In the brain, PA has been shown to lead to changes in cerebral blood flow (CBF) and cerebrovascular reactivity (CVR) (Intzandt et al., 2020). Importantly, PA may slow or even reverse cognitive decline associated with aging (Bherer et al., 2013; Iso-Markku et al., 2024). Furthermore, improvements in cognition resulting from PA engagement correlate with changes in cerebral hemodynamics, including higher CBF (Guiney et al., 2015) and lower cerebral pulsatility (CP) (Mohammadi, Gagnon, et al., 2021). Thus, PA may promote healthy aging in part through maintenance of cerebrovascular function.

The current PA recommendations from the World Health Organisation are identical for adult males and females (Bull et al., 2020) across the lifespan. However, sex differences have been observed in the effects of PA on health, including in the brain. For example, sex differences have been identified in the relationship between grey and white matter volumes and PA among older adults (Intzandt et al., 2023). Moreover, older aged males and females exhibit differences in the association between cardiorespiratory fitness and functional connectivity (Dimech et al., 2019). Although the physiological underpinnings of these sex differences remain unclear, PA appears to offer unique protective mechanisms for each sex (Barha et al., 2019). Therefore, it is likely that males and females benefit in terms of cerebrovascular health from different intensity and dose of PA. Assessing the dose and intensity of PA needed to derive benefit for each sex could furthermore help refine future recommendations in the context of a personalized medicine approach to disease prevention.

Lifelong sex differences alone cannot fully explain the vascular and metabolic changes observed in an aging population; rather, the menopausal transition in females influences these physiological alterations (Newhart, 2013). Menopause has been identified as a significant risk factor for the development of cardiovascular diseases (Anagnostis & Stevenson, 2024). Nevertheless, PA, independent of the menopausal status, has been shown to reduce cardiovascular risk factors (Karvinen et al., 2019). Additionally, menopause is associated with a decline in physical performance, especially strength and power (Bondarev et al., 2021). However, a higher dose of PA can significantly mitigate these reductions in physical performance (Bondarev et al., 2018). Hence, to understand the effects of PA on female cerebrovascular health, it is crucial to stratify according to the menopausal stage.

Magnetic resonance imaging (MRI) can be used to assess several cerebrovascular biomarkers, including CBF, CVR and CP. CVR captures the ability of blood vessels to dilate in response to a vasodilatory stimulus, such as carbon dioxide (CO_2_). CVR is a measure of vascular reserve and has been shown to be more sensitive than CBF to the effects of aging (Gauthier et al., 2013; Lu et al., 2011). The effects of fitness and PA participation on CVR is unclear however, with the extant literature showing both positive (Barnes et al., 2013; Bliss et al., 2023; Tarumi et al., 2015), negative linear (Intzandt et al., 2020; Thomas et al., 2013) and inverted U-shaped relationships (DuBose et al., 2022) in older adults. While some of these effects likely arise from methodological differences, it is currently unknown if some of these contradictory effects may be influenced by sex-specific effects of PA on CVR, or in differential effects of menopause. Here, we will address this gap in the literature. Several methods can be used to measure CVR. Functional MRI (fMRI) blood oxygen level dependent (BOLD) CVR is typically measured as the percent change in BOLD signal in response to a gas manipulation challenge or injection of acetazolamide (Sleight et al., 2021). Recently, a novel technique (Liu et al., 2017) was introduced to acquire CVR exclusively from the BOLD signal during rest, without any respiratory challenge. This method is based on the ability to detect natural end-tidal CO_2_ fluctuations from the lower frequencies of the BOLD signal.

CP measures the extent to which blood vessels expand and recoil in response to fluctuations in blood pressure during the cardiac cycle. CP primarily reflects arterial stiffness through the measurement of pulse pressure waves (Webb et al., 2022). It therefore represents complementary information to CVR, which represents the vasodilatory capacity (Sleight et al., 2021). Higher CP is associated with a decline in microvascular integrity, resulting from increased arterial stiffness (Kim et al., 2021). Sex differences in the effects of PA on CP have been investigated using transcranial doppler (TCD) and 4D flow MRI. Females were shown to exhibit lower CP in the basilar artery with higher levels of PA, whereas males displayed the opposite relationship in the middle cerebral artery. Thus, there may be important sexual dimorphisms in the relationship between PA and CP (Gaynor-Metzinger et al., 2023). MRI studies of CP have typically used 4D flow in large vessels (Morgan et al., 2021). CP at the levels of smaller vessels is also an important marker of cerebrovascular health, though few techniques currently exist to measure it. A technique using near-infrared spectroscopy exists (Mohammadi, Gagnon, et al., 2021), but few MRI-based techniques. Recently, a technique based on hypersampling of the cardiac cycle and the resting state fMRI signal was developed (Voss, 2018), as well as a tool to derive the cardiac cycle from the BOLD signal alone (Aslan et al., 2019). This opens the door to measurement of CP in any dataset that includes a resting state fMRI acquisition. Here, we will assess the effects of PA on CP in sex-disaggregated data, to further understand sex-specific patterns in this relationship.

In summary, the primary objective of this study was to evaluate the sex-specific effects of intensity and amount of time spent in PA on CVR and CP in middle-aged and older adults. The secondary objective was to assess the effect of menopause status in the effects of intensity and dose of PA on CVR and CP. The exploration of these objectives will allow us to better understand how PA participation can be optimized to benefit cerebrovascular health in male and female aging.

## Methods

### Participants

The study included 725 adults (406 females & 319 males) older than 36 years old from the Human Connectome Project – Aging (HCP-A) dataset. The objective of HCP-A is to explore ‘normal’ aging, including vascular risk factors. Participants were screened and excluded for any history of major psychiatric or neurological diseases. More details on the screening process of the HCP-A can be found here (Bookheimer et al., 2019).

From the initial sample, 25 participants (11 females & 14 males) were excluded due PA outliers with PA values above 4 standard deviations, 25 participants (14 females & 11 males) due to missing systolic blood pressure information and 3 participants (3 females) due to missing body mass index information. Individuals older than 85 years old were also excluded due to the variability induced from the superagers profile (25 females & 17 males) (Garo-Pascual et al., 2023). Following the exclusion of these participants, 626 participants (350 females & 276 males) were included in the overall statistical analysis. Additionally, the HCP-A dataset provided demographic information on menopausal status (Harlow et al., 2012). In follow-up analyses on the impact of menopause status, 7 female participants were excluded due to missing menopausal status data. Consequently, 126 premenopausal females and 217 postmenopausal females were included in the statistical analysis.

All participants of the HCP-A dataset provided signed informed consent in accordance with the declaration of Helsinki. Secondary use of this publicly-available data was approved by Concordia University’s Human Research Ethics Committee.

### Physical Activity Measurements

The international physical activity questionnaire (IPAQ) was used to quantify the weekly intensity and dose of PA. The IPAQ was designed to provide a valid measurement of self-reported PA that is reproducible across countries (Lee et al., 2011). The activities from the IPAQ were categorized into metabolic equivalent of task (METs) from the 2011 Compendium of Physical Activities. The PA intensities were classified as walking, moderate (3.0 to 6.0 METs) and vigorous (> 6.0 METs) PA. The total dose of PA was measured as a combination of all PA levels.

The weekly dose of PA in METs were further dissociated into participation levels. Participants were divided into four groups according to the amount of weekly PA participation including: no (0 METs), low (0.01 to 500 METs), medium (500 to 1500 METs) and high (> 1500 METs) PA. These ranges of levels of PA were based on work by (Wang et al., 2019).

### MRI Acquisition

All MRI data were acquired using a Siemens 3T Prisma machine across four sites. The BOLD sequence was obtained using a 2D multiband (MB) gradient echo echo-planar imaging (EPI), providing a 2mm isotropic whole-brain coverage (resolution = 2.0 mm isotropic, repetition time = 0.8s, echo time = 0.037s, MP factor = 8, flip angle = 52°, Volumes = 488). The HCP-A dataset included 4 resting state fMRI runs across two sessions, with two runs per session, each lasting 6.5 minutes. For this study, a randomly chosen single run from the first session, utilizing a posterior-to-anterior phase encoding polarity, was selected. Although the HCP-A dataset included concatenated runs of preprocessed fMRI data, concatenated data was not compatible with the CP measurement and we therefore elected to compute relative CVR and CP from the same fMRI session and encoding polarity. The T1-weighted structural data was acquired using a multi-echo magnetization prepared rapid gradient echo imaging sequence (resolution = 0.8 mm isotropic, echo time = 1.8/3.6/5.4/7.2 ms, repetition time = 2.5s, flip angle = 8°) More details on the HCP-A resting state fMRI acquisition and rationale can be found here (Harms et al., 2018).

### Image Processing

The HCP-A dataset included minimally preprocessed resting state fMRI data, with key preprocessing steps. Freesurfer (Fischl, 2012) and FSL (Jenkinson et al., 2012) toolboxes were employed for both functional and structural data processing (Glasser et al., 2013). Initially, gradient distortion correction was applied to the BOLD images to account for B0 field inhomogeneities. Subsequently, the BOLD time series were aligned to a single band reference to correct for motion artifacts. Echo planar imaging distortion correction was performed using a spin echo field map, further refining spatial accuracy (Andersson et al., 2003). From the preprocessed structural data, the T1-weighted structural image was subsequently registered to the single-band reference (Greve & Fischl, 2009). The resting state fMRI data was non-linearly registered to MNI space using a one-step spline resampling, followed by intensity normalization and brain masking using Freesurfer. The HCP-A preprocessing pipeline, described in detail by (Glasser et al., 2013), was used to ensure consistent and high-quality data processing.

### Relative CVR Analysis

The relative CVR maps were computed from the preprocessed resting state fMRI data. The first step of the relative CVR processing pipeline (***Figure 1***) was to perform smoothing of BOLD images (FWHM = 8mm). A time domain-filtering (0-0.1164Hz) of the BOLD data was conducted with scipy to extract a proxy of the end-tidal CO_2_ (ETCO_2_). This frequency range was chosen based on previous work to isolate the relevant physiological signal associated with ETCO_2_ (Liu et al., 2021). This processing procedure precluded the necessity for a measurement of the ETCO_2_, which is not included in the HCP-A data. Additionally, the filtering of the BOLD signal generates an approximation of resting ETCO_2_ fluctuations, providing relative rather than absolute values. Thus, this CVR method bypasses the need for temporal alignment of the ETCO_2_ data, as the proxy is directly estimated from BOLD signal changes. This contrasts with traditional CVR gas manipulation techniques, where ETCO_2_ is recorded externally at the nose or mouth. The filtered BOLD signal, acting as a proxy of ETCO_2_, was included as a regressor in the general linear model to measure the CVR indices. The CVR index map was normalised to a whole brain CVR average to obtain the relative CVR map. The resting state CVR processing pipeline was based on previous work by (Liu et al., 2017). Relative CVR was averaged over bilateral regions of interests (ROIs) in grey matter of the LPBA40 atlas (Shattuck et al., 2008) normalised to MNI-152 space.

**Figure 1.**
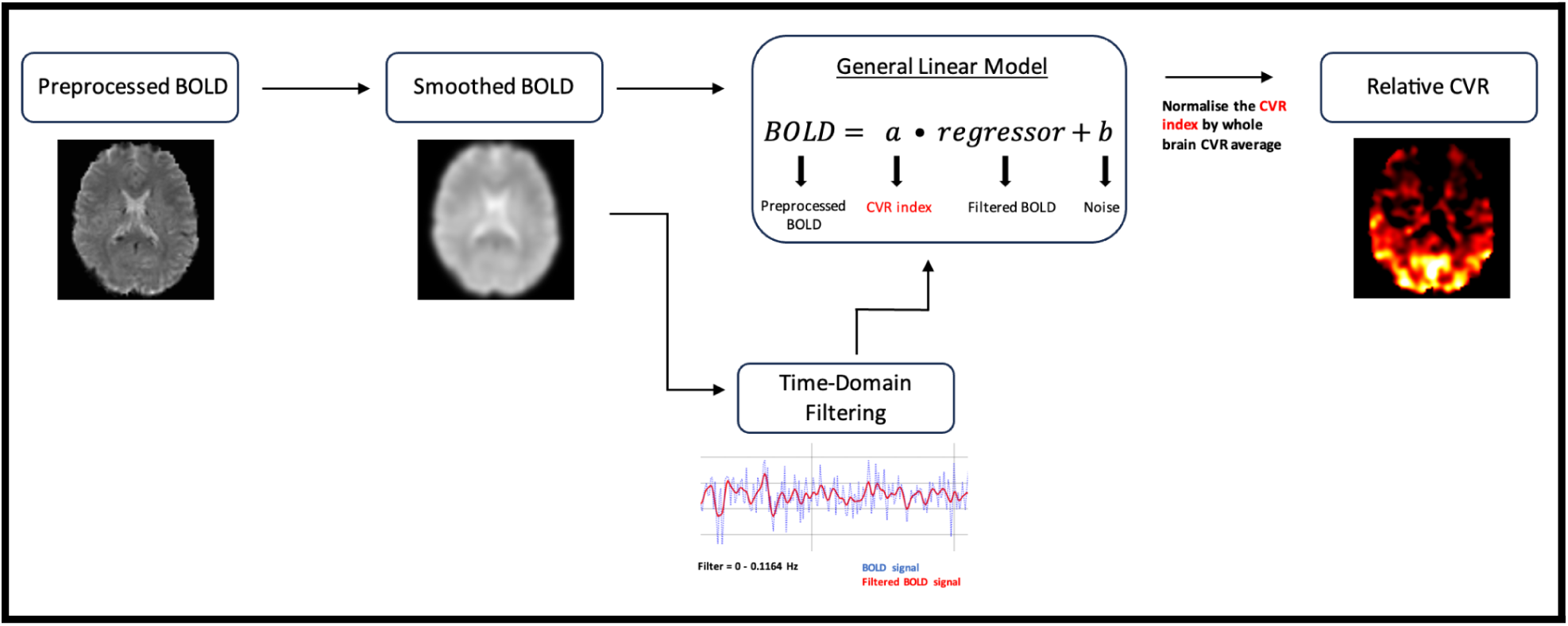
Relative CVR pipeline from preprocessed BOLD data.

### Cerebral Pulsatility Analysis

A proxy of cerebral pulsatility was extracted from the unprocessed BOLD signal using the Hypersampling by Analytic Phase Projection - Yay (Happy) tool (Aslan et al., 2019), a part of the Rapidtide package (doi:10.5281/zenodo.814990). Happy is a Python based tool that can be used to estimate the cardiac waveform from unprocessed resting state fMRI data, and relate this estimate of the cardiac waveform to the voxel-wise BOLD signal timecourse. This technique harnesses the cardiac contamination within the BOLD signal to extract the cardiac cycle. Happy provides a proxy for cerebral pulsatility measured as the BOLD amplitude variations over a cardiac cycle. Happy was derived from Voss and colleagues’ hypersampling technique (Voss, 2018). Because veins are thought to be passively compliant, cardiac variations in the BOLD signal arise as veins passively dilate in response to changes in pressure throughout the cycle. Therefore, while this is not a direct measure of pulsatility of arteries, it is here used as a proxy for local pulsatility.

The unprocessed resting-state fMRI data from the HCP-A was chosen from the same session and phase encoding polarity as the relative CVR analysis to maintain consistency. Cerebral pulsatility (CP) proxy maps were obtained from averaged amplitude BOLD variations over a cardiac cycle using Happy. A brain mask of the fMRI data was extracted using the brain extraction (BET) tool from FSL and was included within the Happy analysis. The CP maps were registered to MNI space using FNIRT from FSL. Additionally, the Jacobian files from the HCP-A were added to the registration of the CP maps to correct from the significant EPI distortions (Glasser et al., 2013). Lastly, a rigid registration was applied with ANTs to improve the alignment of the CP maps to the MNI space. More information on the methodology of Happy can be found here (Aslan et al., 2019). Values were averaged over the bilateral regions of the LBPA40 atlas for subsequent statistical analysis.

### Statistical Analysis

Analyses were performed initially in all subjects with sex as a covariate to investigate sex differences. Then, the analyses were repeated in each sex separately to assess the sex-specific relationships between cerebrovascular properties and PA. This approach is recommended to comprehensively assess the presence of sex-dependent effects (Gigli, 2021; Woodward, 2019). Analyses controlled for age, systolic blood pressure and body mass index. Linear and quadratic multiple regressions were conducted to assess the relationship between the intensity and dose of PA and CVR using ordinary least squares from the statsmodels python program. A Tukey’s Honest Significant Difference Analysis was performed to assess the amount PA participation within a specific intensity or dose of PA on CVR using the pairwise post hoc test from the python library statsmodels. All statistical analyses were repeated to evaluate the relationship between PA and CP. Multiple comparisons across all analyses were corrected with false discovery rate (FDR). Lastly, a corrected p-value threshold of less than 0.05 was used to determine statistical significance.

## Results

The demographic data of all groups are shown in **Table 1**.

**Table 1:**

| Demographic | Males (n = 276) | Females (n = 350) | Pre-menopause (n = 126) | Postmenopause (n = 217) |
| --- | --- | --- | --- | --- |
| Age (years) | 59.7 ± 14.2 | 58.0 ± 13.9 | 44.3 ± 5.8 | 66.4 ± 10.4 |
| SBP (mmHg) | 132.7 ± 16.4 | 128.7 ± 17.6 | 121.8 ± 15.5 | 133.0 ± 17.6 |
| BMI (kg/m<sup>2</sup>) | 27.1 ± 4.0 | 26.9 ± 5.2 | 27.5 ± 5.5 | 26.6 ± 4.9 |
| Total METs (weekly) | 2165.9 ± 1489.4 | 2166.1 ± 1838.9 | 2198.1 ± 1920.2 | 2128.2 ± 1770.8 |
| Walking METs (weekly) | 914.3 ± 1012.5 | 1041.0 ± 1217.7 | 1204.7 ± 1343.8 | 924.7 ± 1064.8 |
| Moderate METs (weekly) | 414.8 ± 481.0 | 504.6 ± 680.5 | 436.7 ± 634.2 | 542.7 ± 704.6 |
| Vigorous METs (weekly) | 837.2 ± 1011.0 | 620.3 ± 914.0 | 556.7 ± 849.9 | 660.5 ± 957.3 |
| Grey Matter Relative CVR | 1.20 ± 0.06 | 1.21 ± 0.06 | 1.22 ± 0.05 | 1.21 ± 0.06 |
|  | Males (n = 267) | Females (n = 337) | Pre-menopause (n = 122) | Postmenopause (n = 208) |
| Grey Matter CP | 201.6 ± 38.7 | 198.9 ± 37.3 | 183.9 ± 34.9 | 208.2 ± 35.8 |

*SBP = Systolic Blood Pressure; BMI = Body Mass Index; METs = Metabolic Equivalent of Tasks; CVR = Cerebrovascular Reactivity; CP = Cerebral Pulsatility.*

### Sex-specific effects in PA and relative CVR

In all participants, a significant negative linear relationship between relative CVR and total PA was observed in the whole gray matter (p=0.010; R=0.241) (Supplementary Figure 1). There was no significant effect of sex on the model. The effects of the different doses of PA on relative CVR in both sexes separately are demonstrated in **Figure 2**. In females, a medium dose of PA (500 to 1500 METs) was associated with higher relative CVR in the cerebral cortex than low (p =0.041; Cohen’s d=-0.45) or high (p=0.007; Cohen’s d=0.40) PA doses. In males, different doses of PA did not lead to significant differences in cerebral cortex CVR.

**Figure 2.**
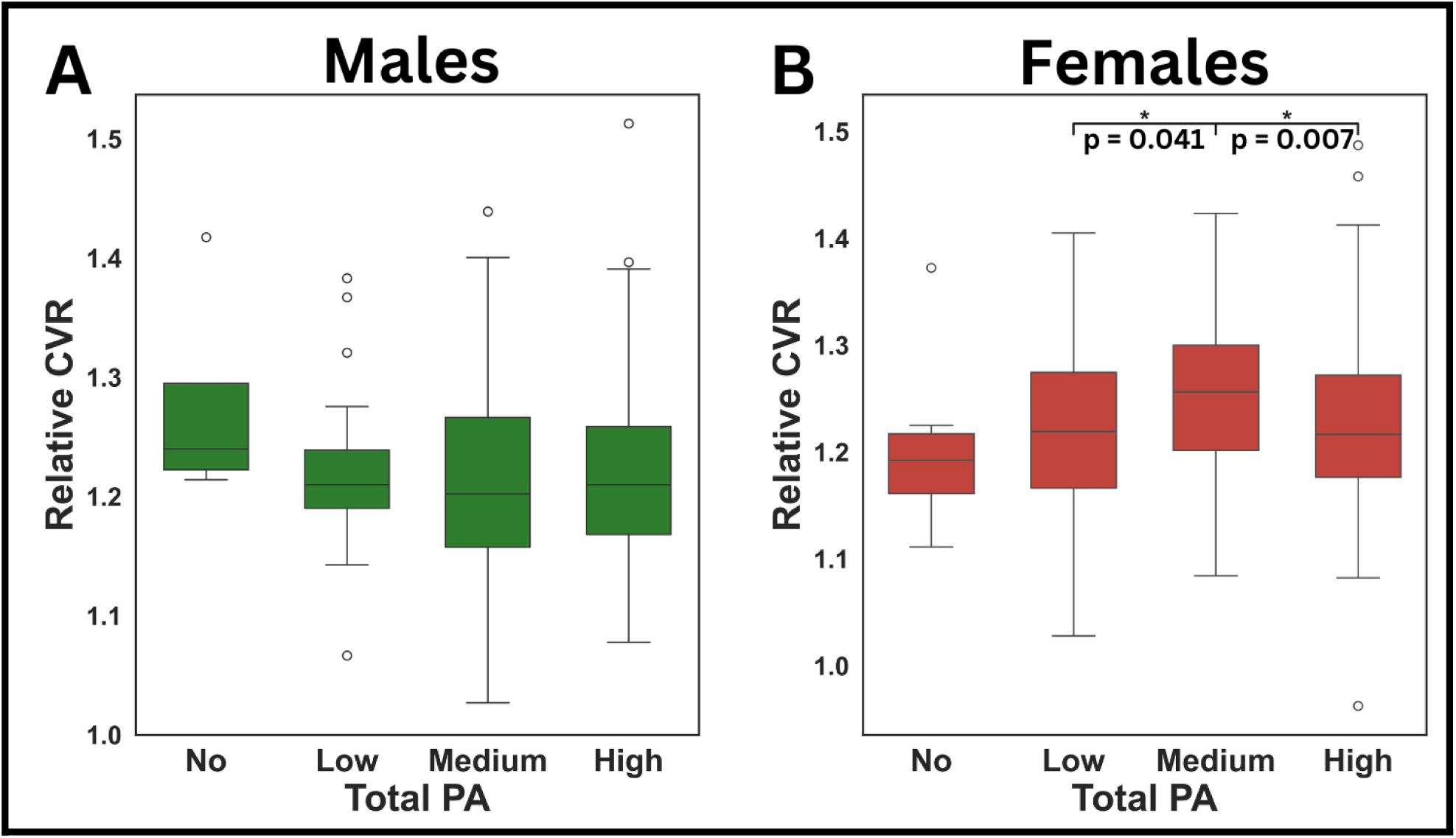
Effects of total PA dose on CVR. *Box plots from Tukey’s tests showing the relationship between relative CVR and total PA dose in males and females. A) Relative cerebral cortex CVR and total PA dose for males (No: n=4; Low: n=28; Medium: n=81; High: n=163); B) Relative cerebral cortex CVR and total PA dose for females (No: n=9; Low: n=48; Medium: n=105; High: n=188). Significance (FDR corrected p < 0.05) is indicated by *.*

### Males

The effects of total PA, as well as dose of specific PA intensities on relative CVR was assessed. Males showed significant negative linear relationships between relative CVR and total PA in the bilateral superior parietal gyri (p = 0.013; R = 0.195) and the insula (p = 0.026; R = 0.158), while a significant positive relationship was observed in the cerebellum (p = 0.035; R = 0.195). Additionally, negative linear relationships were identified between relative CVR and walking PA in the bilateral superior parietal gyri (p = 0.043; R = 0.212) and between relative CVR and vigorous PA in the bilateral cingulate gyri (p = 0.046; R = 0.164). These correlations are displayed in **Figure 3**.

**Figure 3.**
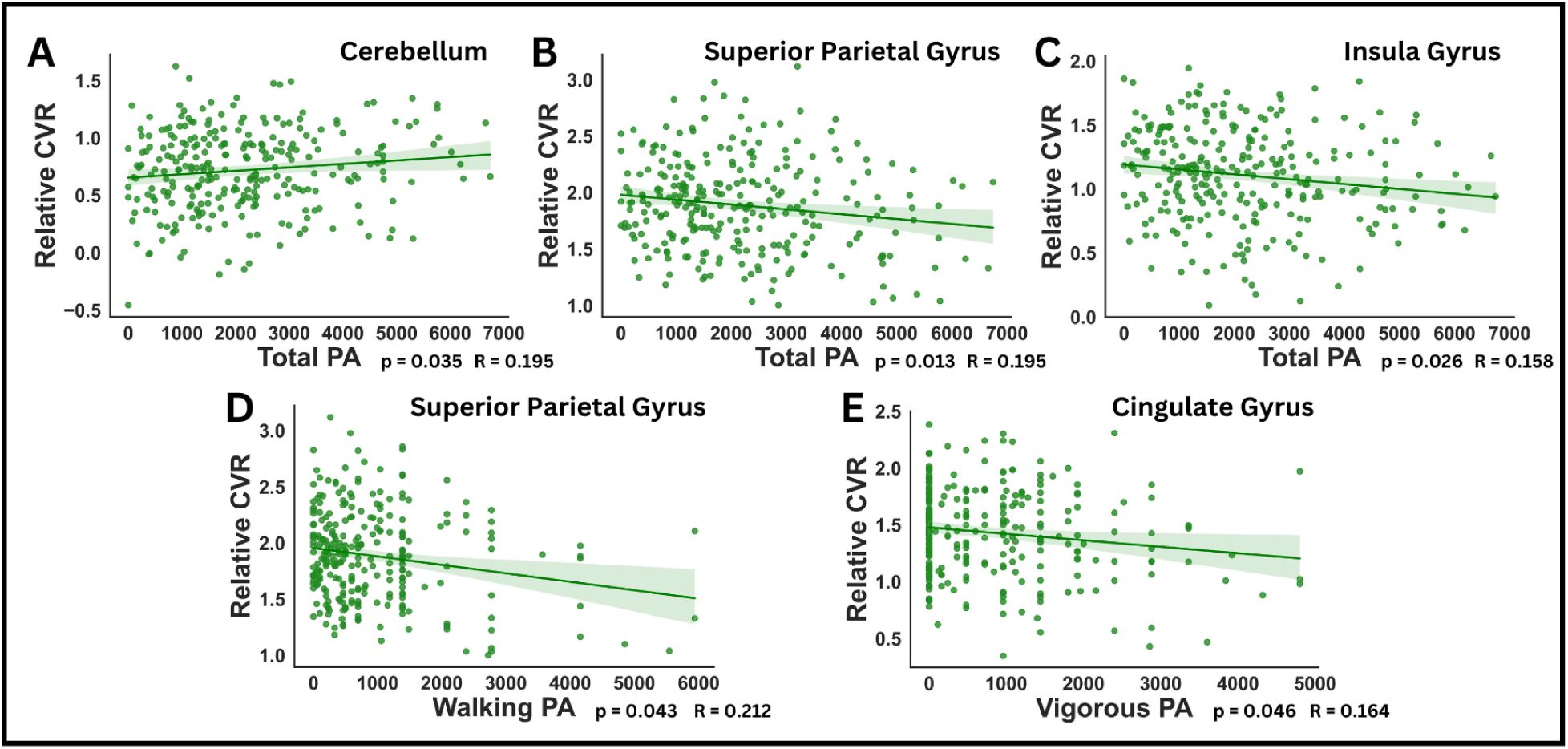
Effects of different PA intensity on CVR in males. *Multiple linear regressions displaying the relationship between relative CVR and different doses of PA at different intensities in males. Linear relations between relative CVR and A) total PA in the cerebellum; B) total PA in the superior parietal gyrus; C) total PA in the insula; D) walking PA in the superior parietal gyrus; E) vigorous PA in the cingulate gyrus. P-values shown were FDR corrected*.

### Females

Females showed a single negative linear relationship between relative CVR and walking PA in the bilateral cingulate gyri (p = 0.033; R = 0.141). Further analyses were conducted with females separated according to menopausal status into premenopausal and postmenopausal groups. Premenopausal females revealed significant negative linear relationships between relative CVR and total PA in the superior occipital gyri (p = 0.026; R = 0.319). We also observed relationships between walking PA in the bilateral superior occipital gyri (p = 0.008; R = 0.361), as well as in the middle occipital gyri (p = 0.011; R = 0.286). In postmenopausal females, we only observed a negative linear relationship with walking PA in the bilateral inferior occipital gyri (p = 0.002; R = 0.237). These results are presented in **Figure 4**.

**Figure 4.**
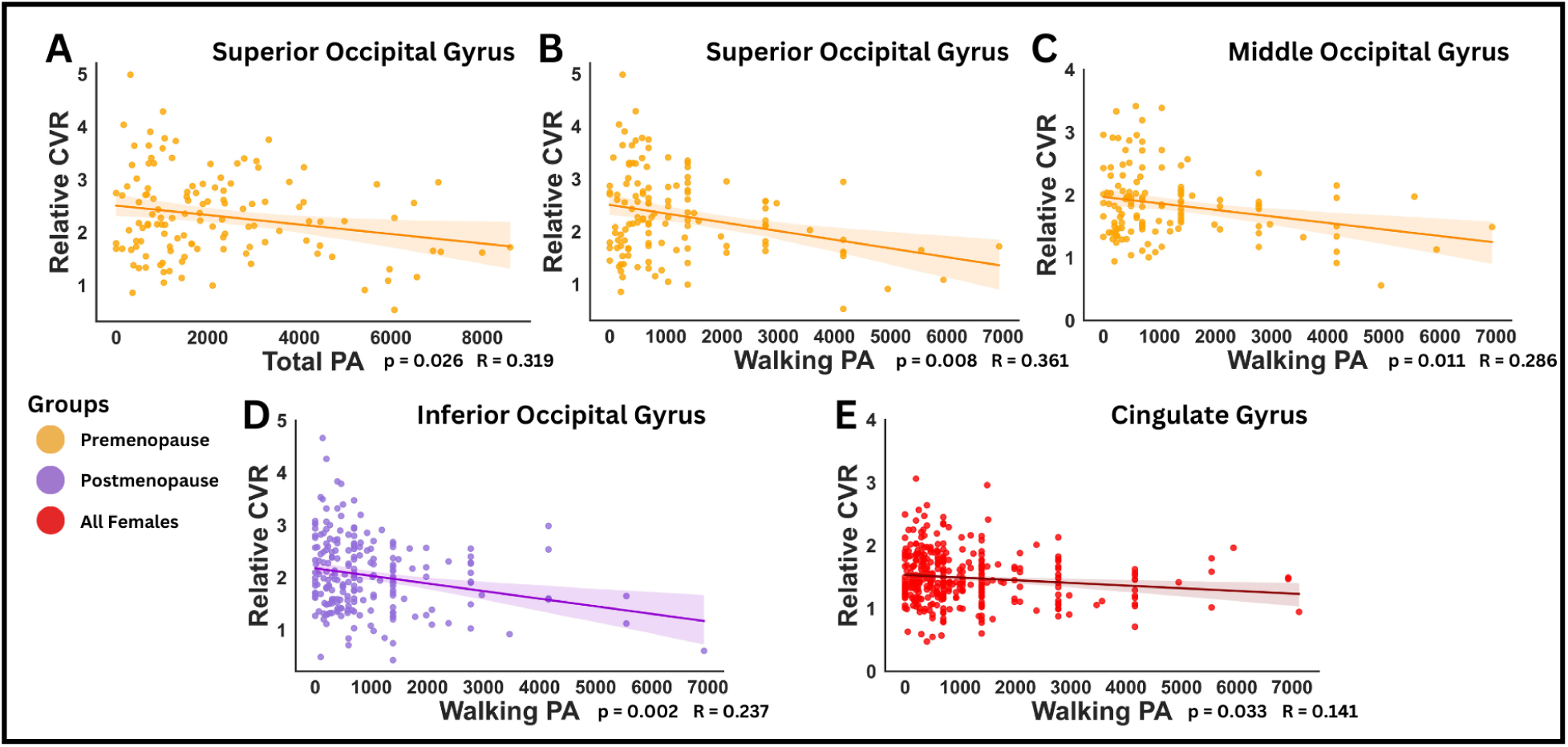
Effects of different PA intensity on CVR in females. *Multiple linear regressions exhibiting the relationship between relative CVR and dose of PA at different intensities in all, premenopausal and postmenopausal females. Linear relations between relative CVR and A) total PA in the superior occipital gyrus; B) walking PA in the superior occipital gyrus; C) walking PA in the middle occipital gyrus in premenopausal females; D) walking PA in the inferior occipital gyrus in postmenopausal females; E) walking PA in the cingulate gyrus in all females. P-values shown were FDR corrected*.

### Sex-specfic effects in PA and CP

There were no significant interactions between PA, sex, and CP when all participants were pooled. The sex-specific effects of total PA on CP are presented in **Figure 5**. Females showed significantly lower CP in medium compared to low PA participation (p = 0.016; Cohen’s d = 0.49) and from high to low PA participation (p = 0.027; Cohen’s d = 0.44) in the cerebral cortex. However, males did not demonstrate any significant effects of the overall dose of PA on CP in the cerebral cortex.

**Figure 5.**
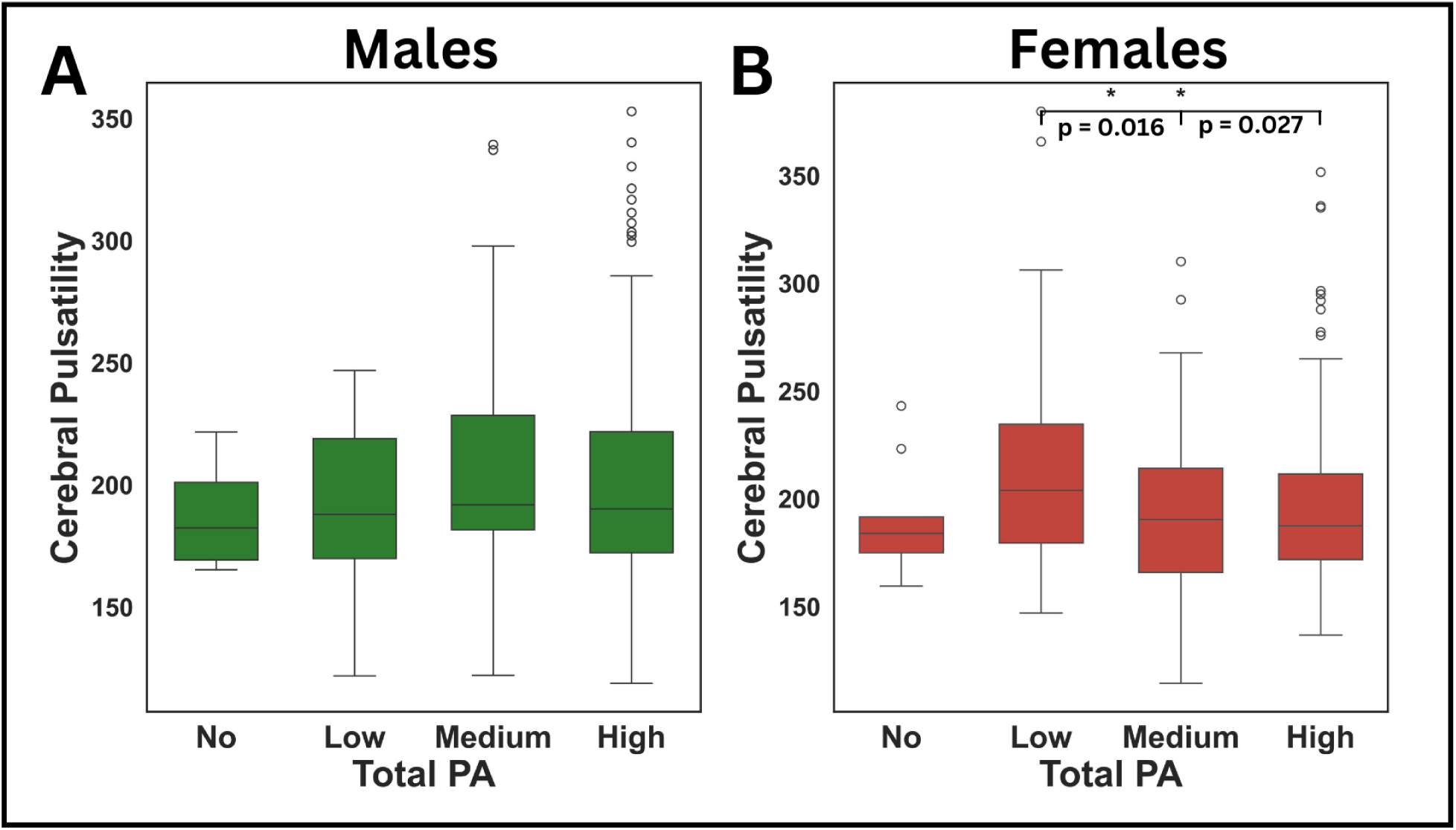
Effects of PA dose on cerebral cortex CP. *Box plots from Tukey’s tests presenting the relationship between CP and total PA in males and females. A) CP and total PA in the cerebral cortex of males (No: n=4; Low: n=27; Medium: n=77; High: n=159); B) CP and total PA in the cerebral cortex of females (No: n=9; Low: n=48; Medium: n=103; High: n=177). Significance (FDR corrected p < 0.05) is indicated by *.*

No effects of PA dose at any intensity was associated with CP in males or when all females were combined in the same group.

The effects of the total PA participation levels on regional CP in all groups are presented in **Figure 6**. In males, a significantly lower CP was observed in the bilateral middle temporal gyri from no to low PA participation (p = 0.049; Cohen’s d = 1.28). In females, the lowest CP was consistently observed at medium total PA dose (p < 0.05). This included a significantly lower CP in medium as compared to low PA participation in all females in the superior temporal gyri (p = 0.039; Cohen’s d = 0.43). Postmenopausal females were the group where this effect was the most pronounced, with lower CP in the no to medium PA participation in the bilateral inferior temporal gyri (p=0.032; Cohen’s d=1.23), cingulate gyri (p=0.039; Cohen’s d=1.37) and the cerebellum (p=0.027; Cohen’s d=1.08), as well as from no to low PA participation in that same region (p=0.028; Cohen’s d=1.05). Conversely, we observed a higher CP from medium to high PA participation in both the bilateral cingulate gyri (p=0.044; Cohen’s d=-0.43) and the hippocampi (p=0.025; Cohen’s d=-0.45).

**Figure 6.**
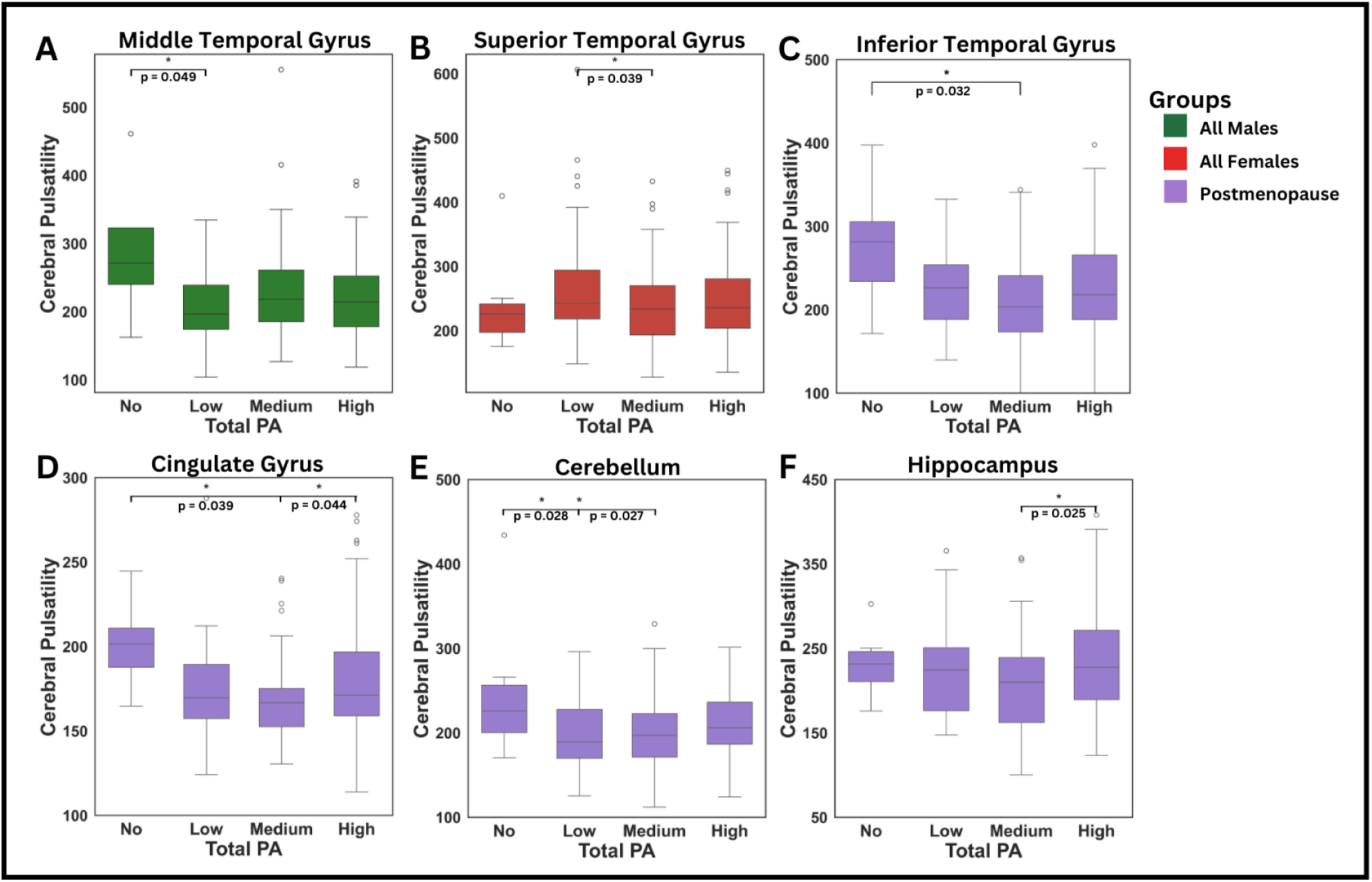
Effects of total PA on regional CP. *Box plots from Tukey’s tests displaying the relationship between CP and dose of PA in males, all females and postmenopausal females. A) Males CP in the middle temporal gyrus (No: n=4; Low: n=27; Medium: n=77; High: n=159); B) Females CP in the superior temporal gyrus (No: n=9; Low: n=48; Medium: n=103; High: n=177); Postmenopausal females CP in C) Inferior Temporal Gyrus; D) Cingulate Gyrus; E) Cerebellum; F) Hippocampus (No: n=6; Low: n=30; Medium: n=59; High: n=113). Significance (FDR corrected p < 0.05) is indicated by *.*

Finally, different PA intensities were not associated with differential effects on CP in males or in the combined group of females. However, some associations were identified once females were segregated into postmenopausal and premenopausal groups. These correlations can be found in (**Supplementary Figure 2)**.

## Discussion

This study assessed sex-specifc effects of dose of PA at different intensities on proxies of cerebrovascular reactivity and cerebral pulsatility. In males, negative linear relationships between relative CVR and PA were shown in several cortical gray matter ROIs particularly for total PA, while a positive linear relationship was identified in the cerebellum. The effects of PA on CP were limited to a single cortical gray matter ROI in males, showing lower pulsatility from no to a low dose of total PA in the middle temporal gyri. In females, relative CVR and CP both showed a pattern whereby the greatest effects of overall dose of PA across all intensities was for a medium PA dose. This medium dose corresponded to the highest relative CVR and the lowest CP values. However, PA dose at specific intensities showed a more complex picture for the relationship between PA and relative CVR, where higher dose of total or walking PA were associated with lower relative CVR, especially in occipital regions as well as the cingulate. This pattern was present in a greater number of regions pre-menopause. In terms of CP, we observed a consistently lower CP for medium level of participation, as compared to no, low and high participation, and this effect was present over more regions in postmenopausal females than other groups. Together, these findings suggest that males and females’ cerebral vasculature responds differently to PA, with menopausal status having a significant effect.

### Sex-specific effects of PA on relative CVR

The results of the CVR-PA relationships indicated predominantly negative linear patterns, with a greater reduction in relative CVR from higher doses of PA. Notably, these findings were both sex-specific and present across several gray matter regions. These regional inverse relationships are consistent with MRI finding showing lower CVR with higher cardiorespiratory fitness or in Master athletes (Intzandt et al., 2020; Thomas et al., 2013). The relationship may be more complex however and this inverse relationship was mostly identified in healthier-than average samples. In contrast, Dubose and colleagues (DuBose et al., 2022) discovered a non-linear relationship between CVR and cardiorespiratory fitness whereby individuals with lower cardiorespiratory fitness showed higher CVR with greater fitness, while individuals at the more highly fit end of the spectrum exhibited a negative relationship. This inverse-U nonlinear effect echoes our findings in the cerebral cortex region in females, where we observed the highest relative CVR in females with a medium dose of total PA, as compared both low or high PA dose. Furthermore, this reduced regional relative CVR in individuals with high PA dose is not unique to females since we observed a similar negative effect in males.

A relationship between PA and relative CVR was also observed for different PA *intensities*. More specifically, males were responsive to all intensities of PA except for moderate PA, whereas all three female groups were primarily sensitive to walking PA. Furthermore, the bilateral cingulate gyri was the only common significant ROI between both sexes. Therefore, the regional variations in the effects of PA on relative CVR between males and females may stem in part from differences in regional perfusion. Consistent with this, sex differences in regional and whole-brain perfusion have been previously reported (Amen et al., 2017). Thus, the effects of PA on relative CVR could be influenced by these sex-specifc regional differences in cerebrovascular changes.

In all groups, the relative CVR-PA relationship was predominantly observed in cortical gray matter ROIs. However, females’ unique relative CVR-PA relationship in the overall region of the cerebral cortex could indicate a greater whole cortical brain sensitivity to relative CVR. It is possible that this relationship is due to the fact that females have higher CBF in the whole-brain, including gray and white matter and greater perfusion differences between gray and white matter (Muer et al., 2024) than males.

Our results highlight an important difference in the PA intensities that are associated with cerebrovascular adaptations. While males displayed relative CVR relationships predominantly with total or vigorous PA, all three female groups (all, premenopausal and postmenopausal females) displayed significant negative linear relative CVR-PA relationships with walking PA. This is in line with studies showing that females benefit more from lower intensities of PA compared to males (Hands et al., 2016). For instance, walking PA has been shown to greatly reduce the prevalence of cardiovascular diseases in females (Bassuk & Manson, 2010). Additionally, females appear to experience improved mental health from lighter PA, while males may benefit more from higher intensities of PA (Asztalos et al., 2010).

### Sex-specific effects of PA on CP

The findings of this study revealed an overall lower CP with higher PA participation across all groups. Since cerebral pulsatility is increased with arterial stiffness, this direction of change is consistent with a beneficial effect of PA on vascular health. Because we did not find any relationship between specific PA intensities and CP, the benefits of PA on CP appear to be primarily influenced by total PA participation rather than a specific PA intensity. These results align with those reported by Mohammadi and colleagues, where a 12-month PA intervention resulted in a significant reduction in CP measured using near-infrared spectroscopy in older adults (Mohammadi, Gagnon, et al., 2021). The improvements in CP from PA are likely due to reduced arterial stiffness which decreases the amplitude of the cardiac pressure wave in the brain (Barnes et al., 2021). Notably, lower CP measured by 4D flow MRI, is strongly correlated with lower arterial stiffness in many large cerebral arteries (Fico et al., 2022). Moreover, greater cardiorespiratory fitness has been inversely associated with 4D flow MRI CP in the internal carotid artery and basilar artery (Maxa et al., 2020). Therefore, total PA, regardless of any specific intensity, likely reduces CP by improving arterial stiffness.

Males showed limited benefits of PA on CP, in contrast to the stronger relationship observed with relative CVR in this group. Therefore, these findings suggest that in males, cerebrovascular health may be more strongly influenced by factors related to vascular reactivity than by protective effects of PA on resting state fMRI-based CP measurement. Consistent with this, the effects of PA on CVR are well documented in males (DuBose et al., 2022; Intzandt et al., 2020; Thomas et al., 2013), while its impact on CP is mainly observed in females (Smith et al., 2021). These sex-specific effects are likely driven by the earlier onset of cerebral artery stiffness in females, resulting from the rapid decline in estrogen levels after menopause, which amplifies cranial pulse waves (Kehmeier & Walker, 2021). Consequently, the effects of PA on CP may be smaller in other groups as compared to postmenopausal females.

Our results in all and postmenopausal females are consistent with previous studies that observed a negative relationship between CP and cardiorespiratory fitness only in females (Lefferts et al., 2022; Zeller et al., 2022). Reduced estrogen after menopause is associated with decreased nitric oxide production and impaired endothelial function (Moreau et al., 2020), leading to greater arterial stiffness. This, in turn, reduces the arteries’ ability to absorb the pulsatility, resulting in increased pulsatility reaching to the brain (Mohammadi, Peng, et al., 2021). Furthermore, older females have been identified as being at higher risk for elevated CP (Lefferts et al., 2020), highlighting the importance of PA at midlife during the transition to menopause and after (Barha & Liu-Ambrose, 2020).

### Potential mechanisms

A few potential mechanisms underlying the negative CVR-PA relationship have been previously proposed, including changes in chemosensitivity (Intzandt et al., 2020; Thomas et al., 2013), cerebral autoregulation and neuroimaging modality differences (DuBose et al., 2022; Intzandt et al., 2020). However, Dubose and colleagues (DuBose et al., 2022) discussed the lack of evidence supporting an attenuation of chemosensitivity due to PA. Additionally, several studies did not report delayed cerebral autoregulation from PA (Aengevaeren et al., 2013; Ichikawa et al., 2013; Perry et al., 2019). Conversely, in individuals with mild traumatic brain injury, PA has been recognised as a potential intervention to improve cerebral autoregulation (C. O. Tan et al., 2014). In contrast, differences in methodology likely explain some contradictory results in the literature. Especially, MRI and TCD measure very different aspects of the vasculature, with TCD being sensitive to changes in blood velocity in large arteries, and MRI measurements being predominantly sensitive to smaller vessels. In fact, MRI and TCD CVR measures have been shown to lack correlation (Burley et al., 2021).

One aspect that may contribute to this negative relationship between PA and CVR is arteriole structural remodelling. The concept of the athlete’s artery was first introduced by Green and colleagues (2012). Endurance trained individuals were shown to have increased lumen size and reduced vessel wall thickness. Evidence from the literature suggests that initially, in untrained individuals, PA may elicit beneficial adaptation to the function of the blood vessel through endothelial improvements, increasing nitric oxide expression (Green et al., 2017) and decreasing oxidative stress (Song et al., 2022). After a few months of PA participation, repetitive bouts of shear stress are believed to lead to permanent vascular structural remodelling (Green et al., 2017). This structural remodelling allows increases in blood flow. Notably, a 16 week aerobic training intervention in young adults has been shown to enhance CBF in association with greater cerebral artery lumen size (Lapidaire et al., 2023). Therefore, vascular remodelling could result in lower CVR by enhancing the capacity to wash out CO_2_ from a greater lumen size. Conversely, Folklow and colleagues discovered that a greater wall-to-lumen ratio induces exaggerated responses to vasodilatory stimuli (Folkow et al., 1958). Thus, low levels of PA may cause an excessive vasodilation response, leading to abnormally high CVR, whereas trained individuals are likely to exhibit a normal CVR response. Our findings revealed, in agreement with Dubose and colleagues (DuBose et al., 2022), that the CVR-PA relationship may not be solely negative, as low to medium levels of PA participation could enhance CVR, at least in the cerebral cortex region in females. Thus, PA may initially improve the function of the cerebral arterioles, increasing CVR, while later structural changes could potentially decrease CVR.

Myogenic tone may also contribute to this negative relationship. Myogenic tone controls how blood vessels contract and dilate in response to changes in blood pressure (Jackson, 2021). Pre-clinical studies have shown that exercise training induces greater myogenic reactivity (Green et al., 2017), meaning more vasoconstriction with increased pressure. In humans, post-exercise recovery has been associated with increased myogenic activity, which improves microcirculation perfusion and facilitates the supply of nutrients to support tissue (Q. Tan et al., 2020). Because CO_2_ increases cardiac output and therefore pressure, higher myogenic tone may lead to a more attenuated response to CO_2_ and a lower CVR. Therefore, PA could potentially enhance the myogenic tone of arterioles, minimising CVR.

Finally, because the relationship between ETCO_2_ and CVR is non-linear, higher resting ETCO_2_ has been linked to reduced CVR (Hou et al., 2020; Tancredi & Hoge, 2013). Individuals with lower respiratory rates have been shown to have elevated ETCO_2_. Moreover, individuals with a higher baseline ETCO_2_ levels have an enhanced cardiorespiratory fitness (Bussotti et al., 2008). Higher body weight increases CO_2_ production which further contributes to greater resting ETCO_2_ levels (Antoine et al., 2018). Additionally, males exhibit higher resting ETCO_2_ levels compared to females regardless of breathing pattern, which may partly explain sex-specifc effects in the CVR-PA relationship (Dhokalia et al., 1998; Kilbride et al., 2003). Thus, lower CVR associated with PA may be influenced by resting ETCO_2_ levels. Overall, these potential mechanisms offer promising insights into beneficial vascular physiological adaptations.

### Limitations

There are a few limitations to this study as the methods to the cerebrovascular measurements rely on resting state fMRI fluctuations. The measurement of relative CVR relies on natural ETCO_2_ fluctuations. For instance, a steady breathing rate diminishes the sensitivity to CVR since the ETCO_2_ changes are small. Therefore, hypoperfused regions may suffer from lower CVR values from a lack of detectible vasodilation. Conversely, individuals with erratic breathing due factors like stress could lead to artificially elevated CVR. The estimation of CP through BOLD originates from venous blood which is not ideal to capture changes in arteries and arterioles. Furthermore, the sensitivity of this CP method is highly reliant on the accuracy of the reconstruction of the heart pulse waveform. Despite these limitations, these techniques are highly valuable, since fMRI BOLD acquisitions are simple to measure and common in both research and clinical routines, making these measurements accessible in already acquired datasets. The fact that these resting state measurements are sensitive not only to the effects of PA, but also to sex specific effects of PA demonstrates their promise as biomarkers of cerebrovascular health.

A second limitation is that PA was quantified from a self-reported questionnaire. Although the IPAQ has been validated across many countries, the results must be interpreted with caution as they are self-reported PA levels. Furthermore, the IPAQ did not assess the participants’ lifetime exposure to PA. Thus, these factors inherently introduced variability in the measurement of PA intensities and doses.

### Conclusions

In summary, this study showed that males and females may benefit differently from varying intensities and doses of PA in relation to cerebral hemodynamics. Females, particularly postmenopausal females, presented more significant effects of total PA on CP compared to males. Because the mechanisms that lead to CVR, CP and their measurement are complex, some of the sex-specifc effects of PA on CVR and CP may be due to different dominant mechanisms of vascular decline and remodelling in males and females. The observed negative relationship between relative CVR and PA may be influenced by physiological mechanisms such as vascular structural remodeling, myogenic reactivity and resting ETCO_2_ levels. Moving forward, research employing gold standard techniques in large, well-characterized cohorts is essential to unravel the complex physiological processes involved and to optimize PA interventions for promoting cerebral vascular health.

## Acknowledgment

Research reported in this publication was supported by the National Institute On Aging of the National Institutes of Health under Award Number U01AG052564 and by funds provided by the McDonnell Center for Systems Neuroscience at Washington University in St. Louis. The HCP-Aging 2.0 Release data used in this report came from DOI: 10.15154/1520707.

Frederick, B, rapidtide [Computer Software] (2016-2024). Available from https://github.com/bbfrederick/rapidtide. doi:10.5281/zenodo.814990

