## Supplemental Figures 1, 2 for "Sex-specific effects of intensity and dose of physical activity on BOLD-fMRI cerebrovascular reactivity and cerebral pulsatility"

**Supplementary**

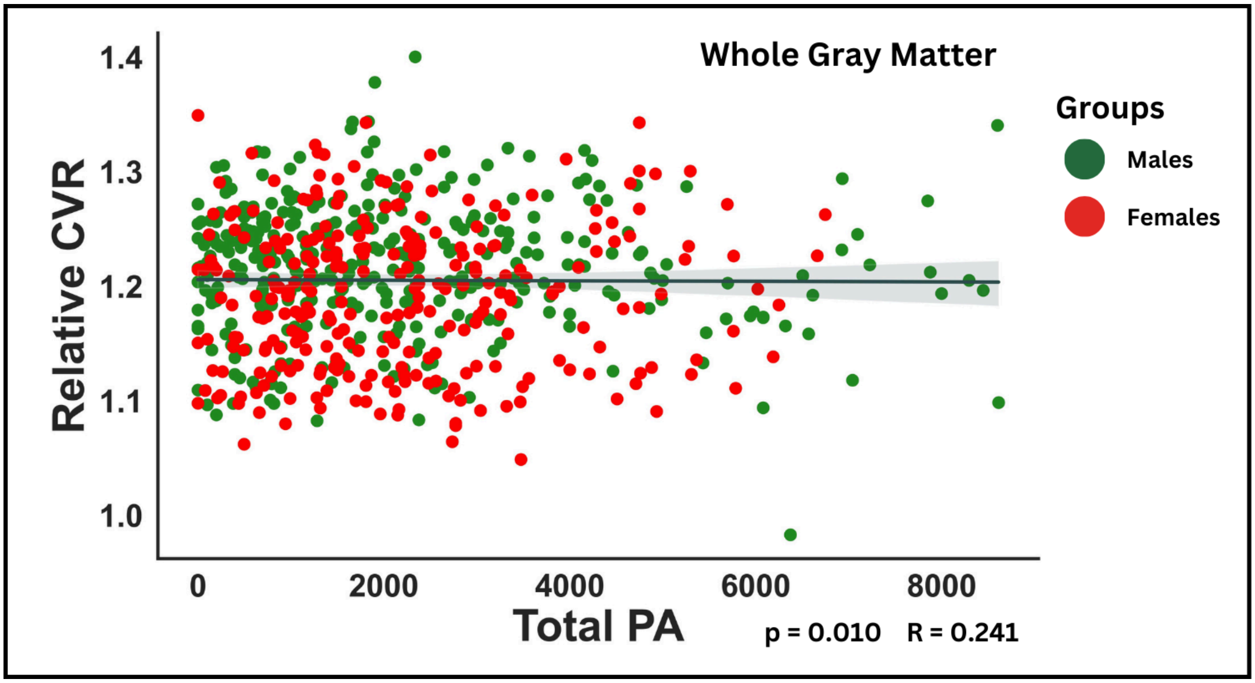


**Figure 1. Effects of total PA on relative CVR in all participants**

*Multiple linear regression showed the relationship between relative CVR and total PA in all participants in the whole gray matter. P-values shown were FDR corrected.*

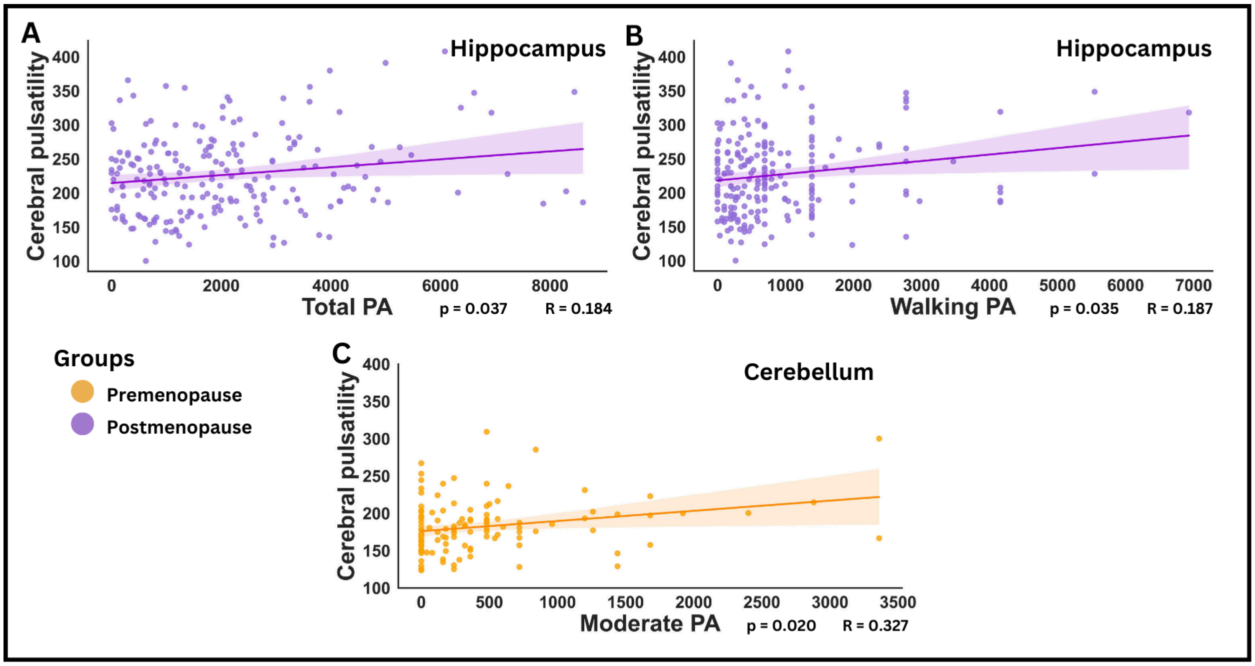


**Figure 2. Effects of different PA intensities on regional CP**

*Multiple linear regressions displayed the relationship between CP and dose of PA at different intensities in premenopausal and postmenopausal females. Linear relations between CP and A) total PA in the hippocampus; B) walking PA in the hippocampus in postmenopausal females; C) moderate PA in the cerebellum in premenopausal females. P-values shown were FDR corrected.*
